# Reconstruction of Gene Regulatory Networks based on Repairing Sparse Low-rank Matrices

**DOI:** 10.1101/012534

**Authors:** Young Hwan Chang, Roel Dobbe, Palak Bhushan, Joe W. Gray, Claire J. Tomlin

## Abstract

With the growth of high-throughput proteomic data, in particular time series gene expression data from various perturbations, a general question that has arisen is how to organize inherently heterogenous data into meaningful structures. Since biological systems such as breast cancer tumors respond differently to various treatments, little is known about exactly how these gene regulatory networks (GRNs) operate under different stimuli. For example, when we apply a drug-induced perturbation to a target protein, we often only know that the dynamic response of the specific protein may be affected. We do not know by how much, how long and even whether this perturbation affects other proteins or not. Challenges due to the lack of such knowledge not only occur in modeling the dynamics of a GRN but also cause bias or uncertainties in identifying parameters or inferring the GRN structure. This paper describes a new algorithm which enables us to estimate bias error due to the effect of perturbations and correctly identify the common graph structure among biased inferred graph structures. To do this, we retrieve common dynamics of the GRN subject to various perturbations. We refer to the task as “repairing” inspired by “image repairing” in computer vision. The method can automatically correctly repair the common graph structure across perturbed GRNs, even without precise information about the effect of the perturbations. We evaluate the method on synthetic data sets and demonstrate advantages over C-regularized graph inference by advancing our understanding of how these networks respond across different targeted therapies. Also, we demonstrate an application to the DREAM data sets and discuss its implications to experiment design.

## 1 Introduction

One of the most exciting trends and important themes in systems biology involves the use of high-throughput measurement data to construct models of complex systems. These approaches are also becoming increasingly important in other areas of biology. For example, mechanistic modeling seeks to describe biomolecular reactions in terms of equations derived from established physical and chemical theory. These theory-driven models use prior knowledge to represent a specific biological system and they work well for pathways in which components and connectivities are relatively well known. While mechanistic modeling approaches should be based on prior biological understanding of the molecular mechanisms involved, when prior knowledge is scarce, a data-driven model may be more appropriate. In such an endeavor, a data-driven model can help us to analyze large data sets by simplifying measurements or by acquiring insight from the data sets, without having to make any assumptions about the underlying mechanism [1].

Among various data-driven modeling approaches, clustering methods are widely used on gene expression data to categorize genes with similar expression profiles. In [2], Eisen *et al.* first demonstrated the potential of clustering to reveal biologically meaningful patterns in microarray data; a cluster analysis for genome-wide expressions data demonstrated that statistical algorithms can be used to arrange genes according to similar patterns and indicate the status of cellular processes. In general, unraveling the complex coherent structure of the dynamics of gene regulatory network (GRN) is the goal of a high-throughput data analysis. Recently, much research has focused on time series gene expression data sets, for example, using functional data analysis techniques for GRN inference [3] [4] [5]. Analyzing these data sets has the advantage of being able to identify dynamic relationships between genes since the spatio-temporal gene expression pattern results from both the GRN structure and integration of regulatory signals. For example, drug-induced perturbation experimental data sets have been combined with temporal profiling which provides the distinct possibility of observing the cellular mechanisms in action [6]. In cancer cells, since signaling networks frequently become compromised, leading to abnormal behaviors and responses to external stimuli, monitoring the change of gene expression patterns over time provides a profoundly different type of information. More specifically, the breast cancer that we study is comprised of distinct subtypes that may respond differently to pathway-targeted therapies [7]. Hence, comparing expression levels in the perturbed system with those in the original system reveals extra information about the underlying network structure. However, since the outcome of data-driven clustering or classification only represents the categorized or clustered responses, they have limitations in inferring the GRN structure directly. As a result, we need extra efforts to infer the network structure from the data.

In mathematical modeling of biological signaling pathways [8] [9] [10], typically only incomplete knowledge of the network structure exists and the system dynamics is known to be sufficiently complex. The challenge has become to show that the identified networks and corresponding mathematical models are enough to adequately represent the underlying system [11]. In the last years, many data-driven inference algorithms have been developed and applied to reconstruct graph structures of GRNs from data. These include Bayesian networks, regression, correlation, mutual information, system-based approaches and *l*_1_-penalized network inference [12] [13] [14] [15] [16] [17] [18] [19] [20]. However, data-driven reconstruction of the network structure itself remains in general a difficult problem; for linear time invariant (LTI) systems, there exist necessary and sufficient conditions for network reconstruction [21]. However, nonlinearities in the system dynamics and measurement noise make this problem even more challenging [22] [23] [24] [25] [26]. In addition, identifying whether important nodes in the graph structure are missing, how many are missing (their multiplicity), and where these nodes are located in the interconnected structure remains a challenging problem.

Until recently, most studies on GRN inference have focused on exploiting a particular data set to identify the graph structure, and have applied the same method to other data sets independently. In addition, although many algorithms use time series gene expression data sets subject to drug-induced perturbations, these perturbations are either assumed to be known [11] or simply ignored. However, such unknown perturbations can cause bias and/or variance in the outcome of the inference algorithm because these unknown perturbations can be considered as corruptions in the measurement and the algorithm is often sensitive to these corruptions. For example, consider a simple inhibition reaction: `A ⊣ B` (i.e., `A` inhibits `B`) and suppose that we perturb `A` and `B` by applying two different inhibition drugs respectively. If the effects of both perturbations are dominant, we may incorrectly infer the relation between `A` and `B` (i.e., we may infer `A → B`) since as `A` decreases, `B` decreases. Thus, in order to infer the GRN correctly, the effect of perturbations should be isolated.

Also, since the effects of the targeted drug can be propagated through the (unknown) underlying network over time, the dynamic responses of gene expressions can be affected directly or indirectly by the drug. For instance, when we design targeted therapies, we obviously know that the response of the target protein is perturbed, so we may assume structured perturbations. However, since these drug-induced perturbations can be propagated and also may have an effect on the other proteins directly, we might only have partial information of these perturbations. Moreover, missing and corrupted data are quite common in biological data sets, and should be properly addressed.

In this paper, we propose a new method to harness various perturbation experimental data sets together, to retrieve commonalities under the sparse low-rank representation, and to improve identifiability of dynamics of GRNs, without any *a priori* information about the GRN structure. Intuitively, without retrieving commonalities, the inferred graph structures from each experimental data set may be biased because each data set has an inherent bias through the perturbation. Thus, the inferred graph structures may not be consistent with each other. By exploiting commonalities across the inferred graph structures, we can estimate bias error due to different perturbations, and correctly identify the common graph structure. We refer to the task as “repairing” inspired by “image repairing” in computer vision. To do this, we first pose the problem as a sparse low-rank representation problem, by formulating the network inference as finding a sparsely connected structure that has low rank over multiple experiments. Inspired by repairing sparse low-rank structure [27] in the computer vision literature, we design a novel convex optimization formulation which enables us to combine temporal data sets from various perturbation experiments. The method can automatically repair the common graph structure from the data sets of perturbed GRNs, even without precise information about the effect of the perturbations. Through numerical examples, we demonstrate the advantage of both dealing with estimation of the perturbation effects and using that information to correctly learn the underlying gene regulatory structure. Also, we demonstrate a possible application using a DREAM data set [28] [29] [30]. We are currently applying this method to biological data sets in HER2 positive breast cancer [6] [7], in which the drugs perturb different parts of the network in each experiment.

The rest of this paper is organized as follows: Section 2 presents the image inpainting method in computer vision by which we inspire the method of repairing common GRNs. In Section 3, we pose the graph inference problem as repairing a sparse low-rank representation and we present the reconstruction of GRN in Section 4. Section 5 presents numerical examples with discussion and Section 6 demonstrates an application of DREAM dataset and discusses the implications and limitations. Finally, conclusions are given in Section 7.

## 2 Background

Although there are deep relationships between clustering and network inference, clustering gene expression data sets and inferring GRNs are tasks usually developed independently. We argue that clustering and network inference can potentially cover each other’s shortcomings since spatio-temporal gene expression patterns result from both the network structure and the integration of regulatory signals through the network [31]. For example, the seminal module networks study [32] and recent study [33] exploit the relationship between clustering and network inference. In [32], Segar *et al.* present a probabilistic method for identifying regulatory modules from gene expression data. Motivated by this, in [33], the authors present a novel approach that infers regulatory programs for individual genes while probabilistically constraining these programs to reveal module-level organization of regulatory networks. In this paper, since we want to reveal the common graph structure of GRN (not limited to module level), we would like to estimate different responses across various perturbations by comparing gene expression levels under the different perturbation conditions and then correctly identifying the common GRN structure.

In Figure 1, we consider collections of time series gene expression of HER2 positive breast cancer cell lines [7] from pathway-targeted therapies involving drug-induced perturbation experiments (`LAP`, `mutant`, `Akti`; for each drug-induced perturbation, we add a single influence for a targeted protein). When a specific protein is perturbed, there are immediate effects on the target protein and compensatory responses on other proteins over time. Thus, comparing gene expression levels in the perturbed system with those in the unperturbed system reveals the extra information about the different cellular mechanisms in action. A dynamical system of the GRN can be modeled as follows:

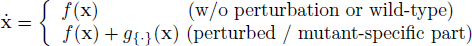

where **x** ∈ ℝ^*n*^ denotes the concentrations of the rate-limiting species, 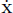 represents the change in concentration of the species, *n* is the number of genes, *f*(·) represents the vector field of the typical dynamical system (or wild-type) and *g*_{·}_(·) represents an additional perturbation or mutant-specific vector field (blue and red edges in Figure 1**A** and **B**). In other words, we have a unified model for wild-type cell line, 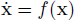 and in the perturbation case, we invoke a single change to the network topology or add a single influence for a specific gene by considering additional vector fields such as *g*_LAP_(·), *g*_AKTi_(·) and *g*_M_(·). Although these additional vector fields affect only a single gene expression at time *t*, their influence can be propagated through the network over time.

**Figure 1.**
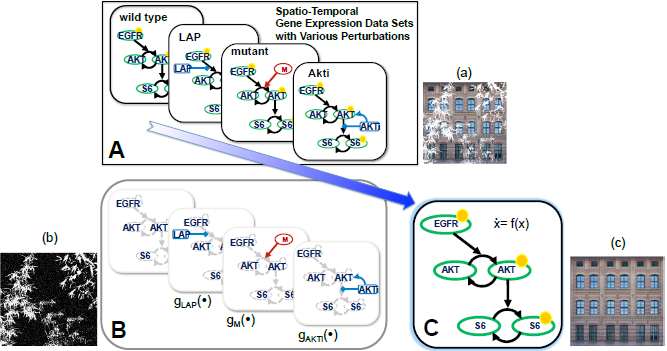
Conceptual diagram of repairing common GRN structure based on collections of time series gene expression from drug-induced perturbation (g_LAP_(`·`), g_M_(`·`), g_AKTi_(`·`)) experiments in HER2 positive breast cancer. In order to show analogous relationship of repairing sparse low-rank texture [27] in computer graphics application, we present each representation with the corresponding illustration such as input image (a), input support (b) and repairing result (c) shown in Figure 2.

By correctly isolating this information in the graph inference problem as shown in Figure 1**B**, we can reduce bias or uncertainties, and hence correctly repair the common graph structure in Figure 1**C**. Intuitively, we can think of these collections of time series gene expression as corrupted graphical images (a) in Figure 1 whose underlying texture is symmetric. By using the properties such as structured regular textures, we can correctly estimate the corrupted region (b) and deal with image completion (c) by repairing the corruption in Figure 1. Our method is inspired by advances in computer vision, which we briefly discuss in the following section.

### 2.1 Overview: Image Inpainting or Completion in Computer Vision

Image completion [34] [35] has been an important but challenging problem widely studied in computer vision and image processing in recent years. The main goal is to automatically recover or regenerate the missing pixel values for a given image with intensity values of certain regions missing (due to corruption). This is inherently an ill-posed problem as the statement of the problem does not ensure there will be a well-defined unique solution, for example, one can fill in arbitrary values for the missing entries [27]. To make this problem well-defined, many researchers seek a solution in which the completed images are in some sense statistically or structurally consistent or follow the same structures as shown in Figure 2.

**Figure 2.**
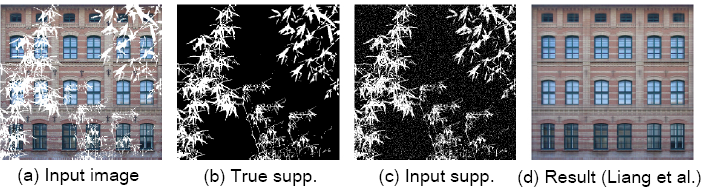
Example of repairing sparse low-rank matrix in computer graphics (Fig. 3 in [27]): This example shows that with 5% support map corrupted, the algorithm recovers the image (d).

There are many methods developed for image inpainting [36] [37] or completion. The first class of methods uses stochastic Partial Differential Equations (PDEs) to describe the structures of the image signals so the remaining completion processes rely on such models and extrapolate the missing part of the image from the given one [38]. In the second class, recent studies [39] [40] perform inpainting based on sparse structures of image patches where patches inside the missing pixel region are synthesized as a sparse linear combination of elements from a patch dictionary. Finally, in the third class, many methods use the patch-based techniques to complete a large missing region or even extend a texture indefinitely relying on restrictive assumptions that the structures of the textures are regular patterns or stationary [27].

### 2.2 Sparse Low-Rank Texture Inpainting and Repairing [27]

In a recent study, Liang *et al.* [27] show how to harness both low-rank and sparse structures in regular or near regular textures for image completion. The method leverages a new optimization formulation for low-rank and sparse signal recovery, with the assumption that the class of images or the structure of the textures has very low intrinsic dimensionality or complexity. More specifically, they consider vectorized images or signals in a high-dimensional space span a very low-dimensional subspace or have a very sparse representation (with respect to a certain basis). They call such texture as “Sparse Low-Rank Texture” and they can correctly repair the global structure of a corrupted texture, even without precise information about the regions to be completed.

Consider a 2D texture as a matrix *I* ∈ ℝ^*m*×*n*^. It is called a low-rank texture if *r* ≪ min(*m*, *n*), where *r* = rank(*I*). For example, all regular, symmetric patterns belong to this class of texture and for such textures, image completion becomes a low-rank matrix completion problem. Suppose *D* ∈ ℝ^*m*×*n*^ is the given image and Ω is the set of observed entries (pixels). The goal is to recover an image *I* of the lowest possible rank that agrees with *D* on all the observed pixels. This completion problem can be formulated as follows:

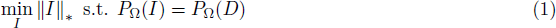

where ‖·‖_*_ is the nuclear norm, convex surrogate of rank of a matrix and *P*_Ω_ is a linear operator that restricts the equality only on the observed entries belonging to Ω. If we want the recovered image to be both low-rank and sparse (in certain transformed domain), the above convex program (1) can be modified as follows:

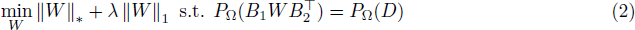

where the matrix *I* can be factorized as 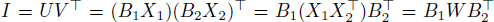 for some orthonormal bases (*B*_1_, *B*_2_) where both *X*_1_ and *X*_2_ will be sparse. Or equivalently, the matrix *W* will be a sparse matrix, which has the same (low) rank as *I*.

Almost all previous methods for image inpainting or completion need information about the support Ω of the corrupted regions. Although this information is manually marked out by the user or detected by other independent methods, in many practical cases, the information about the support of the corrupted regions might not be known or only partially known. Hence, the pixels in the given region Ω can also contain some corruptions that violate the low-rank and sparse structures. In [27], Liang et *al.* model such corruptions as a sparse error term *E*, and modify the above convex program (2):

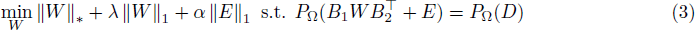

where *λ* and *α* are weighting parameters which trade off the rank, sparsity of the recovered image and the sparse error term. By solving (3), one can decompose the image into a low-rank component and a sparse one as shown in Figure 2. Also, the nonzero entries in the estimated sparse error can help to further refine the support Ω by iterations. In [27], the authors proposed the linearized alternating direction method to solve (3) efficiently.

## 3 Problem Formulation

Inspired by repairing sparse low-rank representation in computer vision, we pose a graph inference problem by formulating the network inference as finding a sparsely connected structure that has low rank over various experiments. Thus, a sparse low-rank representation of GRNs encourages both sparsity of GRNs and commonalities across other’s GRNs.

### 3.1 Formulating Gene Regulatory Networks as a Dynamical System

We consider a dynamical system of GRN described by

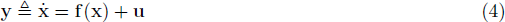

where **x** ∈ ℝ^*n*^ denotes the concentrations of the rate-limiting species which can be measured in experiments; 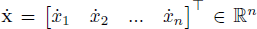 is a vector whose elements are the change in concentrations of the *n* species over time which may not be measured directly in experiments but we could calculate these quantities by interpolating **x** and using numerical derivatives. In [41], the authors pointed out that although data on time derivative can be difficult to obtain especially in the presence of noise, it is possible to estimate the gene expressions relatively accurately by repeating measurement with careful instrumentation and statistics [5] [42]; **f**(·) : ℝ^*n*^ → ℝ^*n*^ represents biochemical reactions, which typically include functions of known form such as product of monomials, monotonically increasing or decreasing Hill functions, simple linear terms and constant terms, since biochemical reactions are typically governed by mass action kinetics, Michaelis-Menten, or Hill kinetics [11] [43]. Since **f**(**x**) determines how the dynamics of 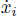 of a protein *i* depends on the expression levels of all proteins, it contains the structural information of the network. **u** ∈ ℝ^*n*^ denotes the control input, for example, drug-induced inhibition or stimulation, for which we only have partial information. For instance, when we inhibit a target protein by drug-induced perturbation, we only know that the dynamics of the targeted gene response may be affected, but we do not know by how large the effect on the dynamics is and how long this effect continues. Moreover, this drug-induced perturbation might also directly affect other proteins in practice.

The nonlinear function **f**(**x**) can be decomposed into a linear sum of scalar basis functions *f*_*b*,*i*_(**x**) ∈ ℝ where we select the set of possible candidate basis functions that capture fundamental biochemical kinetic law [11] [43]:

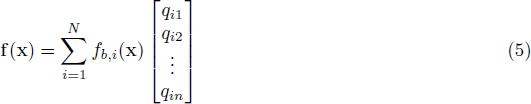

where *N* is the number of possible candidate basis functions and *q*_*ij*_ is the coefficient of the *i*-th basis function for the *j*-th protein response. The biochemical reactions (4) can be written as follows:

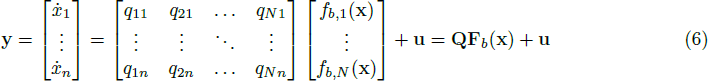

where 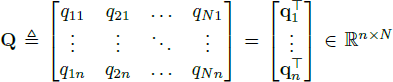, **q**_*i*_ = [*q*_1*i*_ *q*_2*i*_ … *q*_*Ni*_]^⊤^ ∈ ℝ^*N*^ and **F**_*b*_(**x**) = [*f*_*b*,1_(**x**), …, *f*_*b*,*N*_(**x**)]^⊤^ ∈ ℝ^*N*^ is the vector field which includes possible candidate basis functions. Thus, the *i*-th row in **Q** determines the connectivity of the dynamics of the *i*-th protein, through the functional basis in **F**_*b*_. In practice, we can construct **F**_*b*_(**x**) by selecting the most commonly used candidate basis functions to model GRNs, for example, all monomials, binomials, other combinations or Hill function. Thus, any biochemical reactions can be represented by a linear map **Q** and **F**_*b*_(**x**) where **Q** reflects the influence map of GRN structure and **F**_*b*_(**x**) includes all possible candidate functions representing the underlying biochemical reactions. Thus, in order to infer the graph structure, we want to recover **Q** from the measured response **x**, 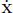 with the chosen basis functions **F**_*b*_(**x**).

#### Example 1

*Consider simple nonlinear ordinary differential equations (ODEs) without drug-induced perturbation:*

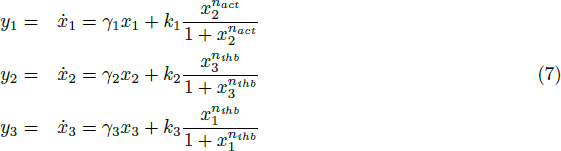

*where x*_*i*_ *denotes the concentration of the i-th species*, 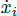 *is the change in concentration of the i-th species, the γ*_*i*_ *denotes protein decay rate, the k*_*i*_ *denotes the maximum promoter/inhibitor strength. Here, there is no perturbation input* (**u** = **0**). *Also, n*_*act*_ > 0 *represents positively cooperative binding (activation) and n*_*ihb*_ < 0 *represents negative cooperative binding (inhibition). The set of ODEs corresponds to a topology where gene 1 is activated by gene 2, gene 2 is inhibited by gene 3, and gene 3 is inhibited by gene 1 as shown in Figure 3(left). We can write (7) as follows:*

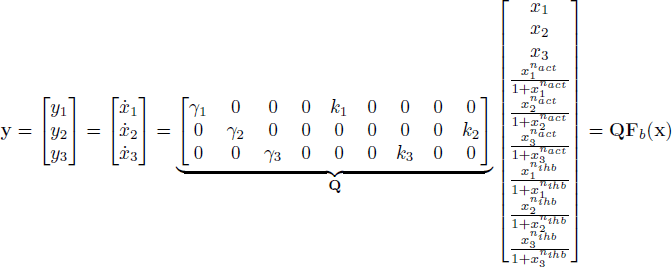

*where* **Q** ∈ ℝ^3×9^ *represents the influence map. We can also consider drug-induced perturbation or control input* **u** *as shown in Figure 3(right):*

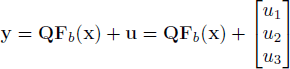

By formulating the dynamics of GRN into Equation (6), we are able to extract the graph structure of GRN into **Q**. To adopt the methods in repairing sparse low-rank representation to our problem, we need to form a matrix representation of the unknown network structure, using the form of matrix *W* in Equation (3). Thus, in the next section we organize the dynamic equations of the GRN into the desired matrix form amenable for repairing sparse low-rank representation.

**Figure 3.**
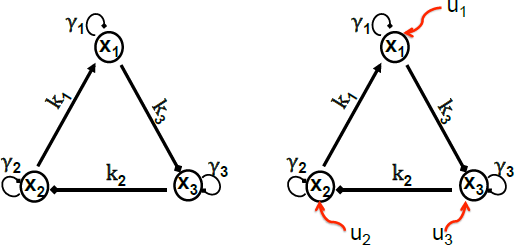
A graph representation of nonlinear ODEs (Example 1, *n* = 3, *N* = 9 and **Q** ∈ ℝ^3 × 9^): among 27(= 3 × 9) components, only 6 components are non-zero and they represent graph edges, (left) without perturbation (right) with perturbation **u**.

### 3.2 Organizing GRN Dynamic Equations into Sparse Low-rank Representation

Suppose the time series data are sampled from a real experimental system at discrete time points *t*_*j*_. By taking the transpose of equation (6) and vectorizing **Q** as **s**, we obtain

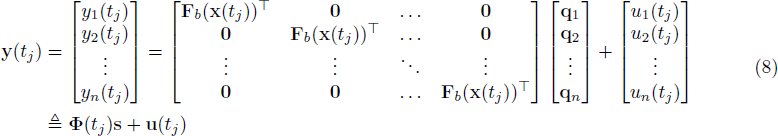

where **y**(*t*_*j*_) ∈ ℝ^*n*^, Φ(*t*_*j*_)^1^ ≜ (**F**_*b*_(**x**)^⊤^ ⊗ **I**_*n*_) ∈ ℝ^*n*×*N*·*n*^ (⊗ denotes Kronecker product) and **u**(*t*_*j*_) ∈ ℝ^*n*^. Note that we define 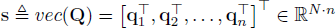 by vectorizing **Q**.

Since we have partial information about **u**(*t*_*j*_), we want to exploit this information to reconstruct **s**. If we inhibit the *k*-th gene by a drug, for example, we know that

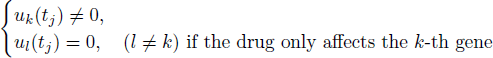

From Equation (8), we expect that the *k*-th component of **y**(*t*_*j*_) is corrupted due to unknown perturbation *u*_*k*_(*t*_*j*_). If we use this corrupted data to reconstruct **s**, the estimated **q**_*k*_ corresponding to this corruption (*u*_*k*_(*t*_*j*_) ≠ 0) might be biased. However, we can simply consider the corrupted part as unmeasured and ignore it for reconstruction, possibly using another experimental data set for this part. Since we consider various perturbation experimental data sets, we denote Equation (8) as follows:

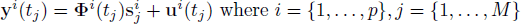

where the superscript *i* denotes the *i*-th experiment and the subscript *j* denotes the time step for each *i*-th experiment, and we have *p* different experiments and *M* time steps for each experiment. Then, we define 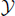, 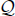 and 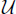 as follows:

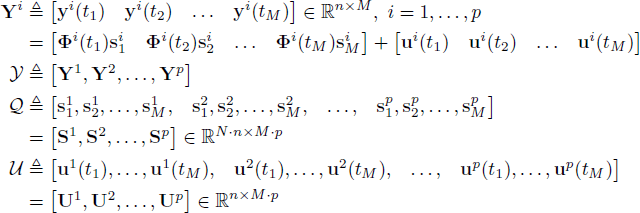

where **Y**^*i*^ represents the measured dynamic responses, 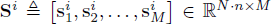 represents the unknown GRNs structure for the *i*-th experiment over time and **U**^*i*^ ∈ ℝ^*n*×*M*^ represents partially known perturbation input over time for the *i*-th experiment.

#### Example 2

*(Recall* **Example 1***) we can denote* **s** *as follows:*

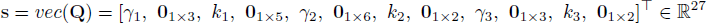

*Among 27 components, only 6 nonzero components represent edges of the graph structure as shown in Figure 4. Also, Figure 4(right) represents a sparse low-rank matrix representation of* 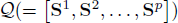 *where each parameter of* **S**^*i*^ *may change over time or across experiments but* **S**^*i*^ *or* 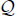 *is structurally consistent or follows the common graph structures. Thus, our goal is finding a sparsely connected structure that has low rank over various experiments.*

**Figure 4.**
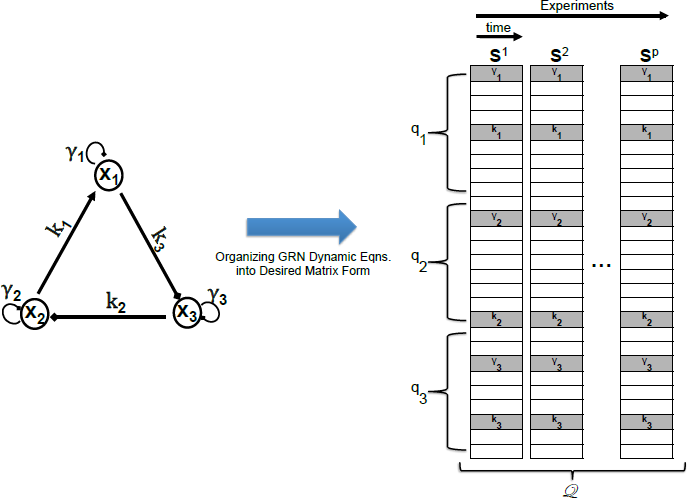
Example 2: A sparse low-rank matrix representation of 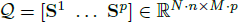 from nonlinear ODEs (Example 1) where each gray box represents nonzero component in **S**^*i*^ (i.e., graph edges) and each white box represents zero components (i.e., no edge).

Without loss of generality, since the GRN structure is assumed to be sparse and not to change over time, there are many zero rows in **S**^*i*^, and hence there are many zero rows in 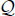 as shown in Figure 4. For example, if parameters of the influence map do not change over time, **S**^*i*^ ∈ ℝ^*N*·*n*×*M*^ can be represented by 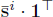 (i.e., **S**^*i*^ has rank 1 where 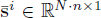 is assumed to be sparse and **1**^⊤^ ∈ ℝ^1×*M*^). Moreover, although all treatments result in down-regulation or up-regulation of gene regulatory signals, they can be well represented by Φ^*i*^(·) and the topology of the underlying influence map may not be changed. Therefore, 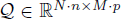 can be well represented by a matrix, which is both sparse and low rank. For instance, if the underlying graph structure is *r*-sparse, then 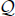 can be represented by 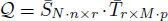 where *r* ≪ min(*N* · *n*, *M* · *p*).

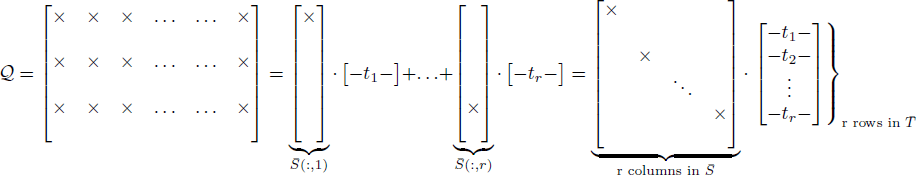

where × denotes non-zero components representing edges in the graph structure (these are shown in gray box in Figure 4(right)), 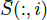 represents the *i*-th column of 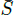 and *t*_*i*_ ∈ ℝ^1 × *M* · *p*^ is the *i*-th row vector of 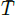 representing variation over time and across experiments. For 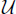, we have (partial) information on the structure of the corrupted region (initial support Ω in Equation (1)) since we only have information about drug perturbations, even without precise information about these effects.

By formulating a dynamical system as a GRN, we can construct a sparse low-rank matrix 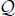, which enable us to use the sparse low-rank texture repairing method in Equation (3) to recover 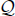. In this paper we make the following assumptions:

#### Assumption 1

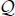 *can be represented by a matrix of sparse and low rank. More precisely, parameters of the influence map are assumed to be fixed over time for each experiment.*

#### Assumption 2

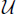 *is (partially) block-sparse and these nonzero blocks can be distributed uniformly by designing experiments. Also, we have partial information about, the position of these blocks.*

Assumption 1 asserts that 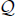 can be represented by a sparse low-rank matrix, so that we can correctly repair the common graph structure from various perturbation experimental data sets. Without loss of generality, since GRNs are assumed to be sparse and 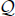 denotes the underlying GRNs over time and across different experiments, the structure of 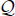 has very low intrinsic dimensionality (sparse low-rank). Assumption 2 then states that our perturbations should excite the network uniformly, in order to retrieve the common structural and temporal information from which we can correctly repair the common GRN structure. Intuitively, if we corrupt or block entire rows of the image in Figure 2(a), there is no way to correctly repair these rows. Similarly, if the responses of a specific protein are always corrupted directly by drug-induced perturbation across the entire experiments, there is no way to repair the corresponding structure.

#### Example 3

*Consider a simple linear system* 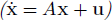 *where n* = 3:

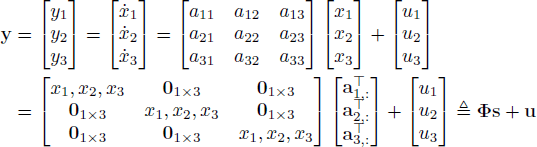

*where* **a**_*i*,:_ *represents the i-th row of A*, Φ ∈ ℝ^3×9^, **s** ∈ ℝ^9^ *and* **u** ∈ ℝ^3^. *Consider the time series data which are sampled at discrete time points (M* = 4):

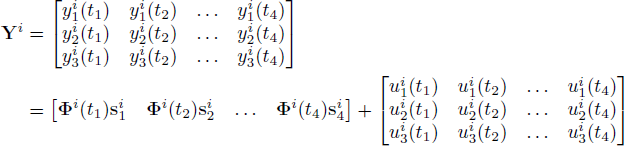

*For the first experiment* **Y**^1^*, suppose that we perturb node* 2.

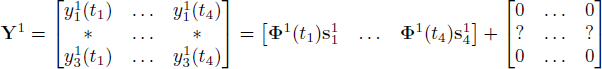

*where* ? *denotes nonzero but unknown input component and* * *represents certainly corrupted responses in y followed by (8). Without loss of generality, since the drug induced perturbation can be propagated through the underlying regulatory network over time and the drug may have (unknown) effects on other proteins, the region of corrupted responses in y may be large (i.e., there may be many* * *in* **Y**^1^).

*Similarly, suppose that for the second experiment* **Y**^2^ *we perturb node* 3, *and for the third experiment* **Y**^3^ *we perturb node* 1. *Then we can construct the matrices as follows:*

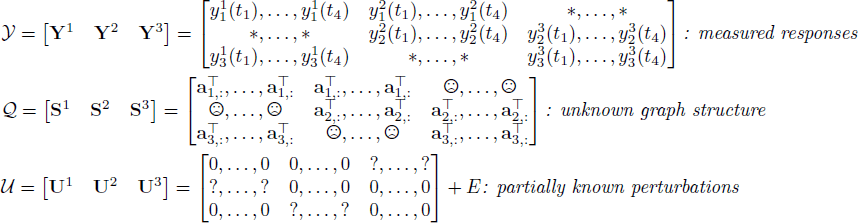

*where* 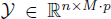, 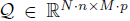, 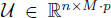 *and E* ∈ ℝ^*n*×*M*·*p*^ *represents additional unknown corruption, for example, drug may affect other protein directly or there may be missing and corrupted data. Thus, E represents such corruptions as a sparse error term shown in Equation (3). Also, for reconstruction of* 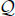*, we may consider* * *as unmeasured part due to corruption in y (denoted by* \**) and ignore it for reconstruction. Thus, we repair* 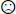 *from the inferred GRN possibly using another experimental data set for this part.*

Figure 5 illustrates the overall procedure; Figure 5(left) represents a schematic overview of the drug-induced perturbation responses over time on the GRN. This schematic overview summarizes the immediate effects of drug-induced perturbations and compensatory responses over time. For example, perturbations (blue arrow) have the immediate effects on up/down (positive/negative) regulation of signaling at the immediate target and other proteins (these are shown in green). Unperturbed dynamic responses are shown in red and non-triggered dynamic responses are shown in gray. For example, a high concentration of a drug causes consequent shut-down of regulatory network. Hence, without handling drug-induced perturbation properly, the result of any inference algorithms may be biased. For instance, if the GRN is only partially triggered, there is no way to infer the entire network structure using the corresponding data set.

**Figure 5.**
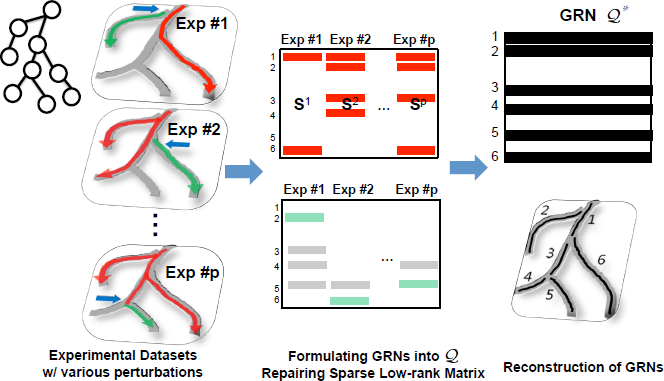
Conceptual diagram of overall procedures: (left) schematic overview of the drug-induced perturbation responses over time with GRN structure (shown in gray), (middle) formulating GRNs into a matrix form and repairing sparse low-rank matrix and (right) the common GRN structure.

Figure 5(middle) illustrates matrix 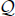 where we separate them as uncorrupted (middle, top) and corrupted components (middle, bottom) respectively with respect to drug-induced perturbations. Similar to Figure 2, the matrix 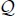 is corrupted by various perturbations. Since we have partial information of these perturbations 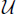 and the 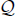 follows the common graph structure of GRNs, we can recover the common GRN structure by correctly repairing the corrupted structure of GRNs as shown in Figure 5(right), even without precise information about the corrupted regions and values to be completed.

## 4 Reconstruction of GRNs via Repairing 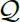

Since we construct the desired matrix form in the previous section, we will show how to harness both sparse and low-rank structure for inferring the common graph structure from various perturbation experimental data sets.

### 4.1 Repairing 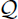 by Refining Support Estimation

Similar to Equation (3), we consider the following optimization problem:

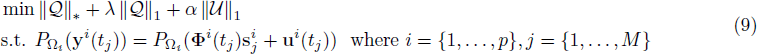

where *λ* and *α* are weighting parameters which trade off the rank and sparsity of the recovered graph structure, and the influence of the drug-induced perturbation respectively. In practice, we can use these parameters as tuning parameters to extract meaningful graph structure and recover the common graph structure which can be represented by sparse low-rank matrix 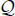 from various drug-induced perturbation data sets. For example, if we want to find the common core structure of GRN, we may set both *λ* and *α* small values in order to penalize the commonalities (i.e., rank properties). Thus, by adjusting these parameters, we can also narrow down the key components of GRN structure. Also, here we define a linear operator 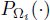 that restricts the equality only on the entries belong to Ω_*i*_ and we could consider a simple set in ℝ^*n*^ [27]:

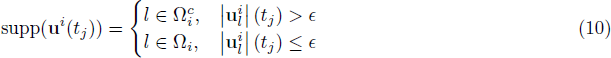

for some threshold *ϵ* > 0. Thus, 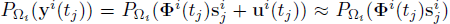. Also, we could estimate the support of **u**^*i*^ using a more sophisticated model to encourage additional structures such as spatial or temporal continuity [27] or to incorporate *a priori* information such as positive perturbation 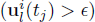 or negative perturbation 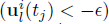.

Since we have partial information about **u**^*i*^(*t*_*j*_), we only use the uncorrupted information to reconstruct the graph structure. For the corrupted part, we estimate the corruption signal 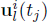, update set Ω_*i*_, and solve the optimization iteratively. We could iterate between the reconstruction and refine the support estimation as follows:

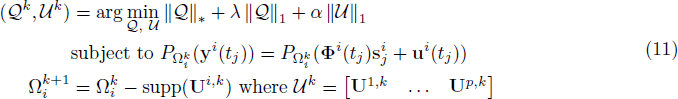

where superscript *k* represents the iteration step. We could iterate the above procedure (11) till it converges and then we can recover the optimal 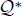 and estimate the corresponding 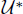.

### 4.2 Handling a Large Number of Candidate Basis Functions

In computer graphics applications, although being low-rank is a necessary condition for most regular, structured images, it is certainly not sufficient [27]. In order to repair a more realistic regular or near regular pattern (typically piecewise smooth), Liang *et al.* consider additional structures by introducing certain transformed domains [27]. In the biological setting, since we select the set of possible candidate basis functions that capture fundamental biochemical kinetics and the number of sample time steps (*M*) is limited in biological data sets, the number of rows in 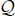 is easily greater than the number of columns. Thus, 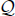 becomes a tall matrix which easily has full column rank. Since we want to encourage the common graph structure across column spaces, being low-rank may not be sufficient to repair the same structure, especially considering a large number of basis functions with limited time samples. For example, when we consider a tall matrix, reducing rank of 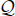 may not encourage the common graph structure across the different experiments; There could be variations across the horizontal direction without affecting the rank or sparsity of the matrix. Hence, in order to recover a more realistic regular or near regular pattern across column space of tall matrix 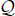, we modify the above convex program in Equation (9) as follows:

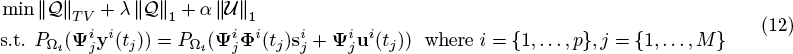

where instead of the nuclear norm 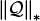, we minimize the total variation 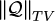 defined by:

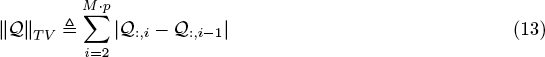

where 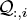 denotes the *i*-th column of 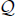. Also, compared to image repairing, we have extra information such as the time invariance of the GRN structure and thus we can impose additional constraints. For example, parameters of the influence map for each experiment *i* are assumed to be fixed over time for each experiment (i.e., 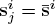) followed by Assumption 1. Thus, we regularize the variation across column spaces in 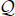 in order to recover meaningful common GRN structure; Otherwise, for example, if we use 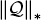 with a large number of basis functions, it is often hard to repair common or near common graph structure.

Lastly, we introduce transformations 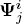 in Equation (12) by which the components of the sensing matrix **Φ**^*i*^(*t*_*j*_) can be made more uniformly distributed so that we reduce the coherence and improve identifiability, as discussed in [11]. Here, we simply use a randomly chosen matrix for 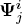. Since randomly chosen matrices spread out the component of **Φ**^*i*^(*t*_*j*_) and **u**^*i*^(*t*_*j*_) uniformly, it helps to differentiate the influence from highly correlated bases in **Φ**^*i*^(*t*_*j*_) in practice.

## 5 Numerical Examples

Here, we consider synthetic experimental data sets for both linear and nonlinear systems.

### Example 4

*Consider a simple linear system with n* = *N* = 10, *M* = 6 *and p* = 10. *Figure 6(a) shows a randomly generated graph structure which has* 10 *nodes and* 23 *edges. We perturb this system with different inputs and measure the response x and* 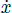. *By repairing the sparse low-rank matrix* 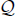*, we reconstruct the GRN in Figure 6(b)***B**. *Also, we compare with the L*1 *result in Figure 6(b)***C** *as follows:*

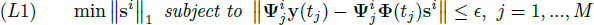

*where ε is constant and* 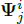 *denotes randomly chosen matrix to reduce coherence. Note that if there is no drug-induced perturbation or the exact* **u**^*i*^(*t*_*j*_) *is assumed to be known [11], we can subtract* **u**^*i*^(*t*_*j*_) *in from* **y**(*t*_*j*_) *and then we can set ϵ* = 0 *(equality constraint). Although we do not know the exact* **u**^*i*^(*t*_*j*_) *in practice, many existing inference methods simply ignore this effect (i.e., without handling* **u**^*i*^(*t*_*j*_) *properly) and thus the inference result can be biased. Since L*1 *uses the corrupted response and identifies the underlying graph structure for each data set independently, the reconstructed results show inconsistency across different perturbation experiments in Figure 6(b)*C. *Thus, drug-induced perturbations cause bias or variance in the result of the L*1-*inference method which is often sensitive to these corruptions. However, since the proposed method incorporates all information across different experimental data sets, we can estimate the corrupted input in Figure 6(b)*F *and reconstruct the GRN via repairing the common structure. Also, although we only use partial information of perturbations in Figure 6(b)*E *at the initial iteration, we can estimate the corrupted input fairly well in Figure 6(b)*F.

**Figure 6.**
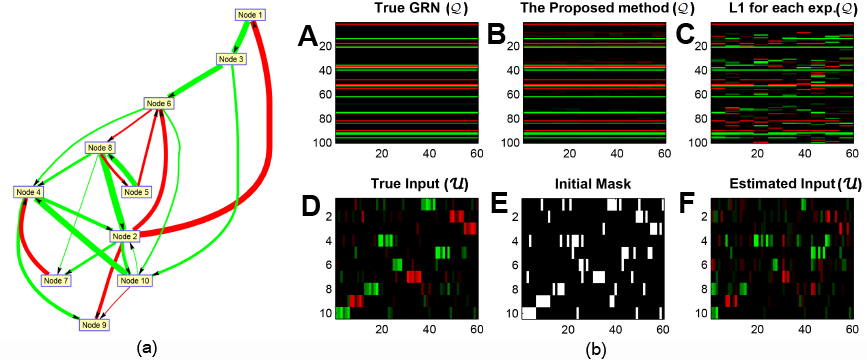
Example 4: (a) GRN where red represents activation edge and green represents inhibition edge (b) recovery of GRNs via repairing sparse low-rank matrix where we compare with *L*1 optimization where red represents an activation edge, green represents an inhibition edge and black represents no edges. **A** represents the true GRN, **B** represents the inferred GRN by the proposed method, **C** represents the inferred GRN by *L*1 optimization, **D** represents the true corruption, **E** represents the initial mask and **F** represents the estimated corruption. Since we consider a linear system with *n* = *N* = 10, there exist *n* · *N*(= 100) possible edges. Here, 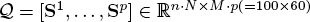 and 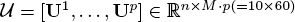 where *M* = 6 and *p* = 10.

### Example 5

*(Increasing p, the number of perturbation experiments) Consider a linear system in Example 4 with increasing p* = 1, …, 10. *Figure 7 shows the reconstruction result with respect to p. Figure 7(a) shows the reconstruction result and (b) shows mean absolute error with error bars for L*1 *and the proposed method. As p increases, since we have a more chances to perturb the underlying system uniformly well, the proposed method can not only reduce reconstruction errors (solid line) but also reduce variance (error bar). In practice, since we do not know the underlying structure, reducing uncertainties is quite important to provide us more convincing reconstruction results. L*1 *reconstruction results show that the reconstructed graph structure is only partially consistent with the true structure and has large uncertainties in Figure 7(b).*

**Figure 7.**
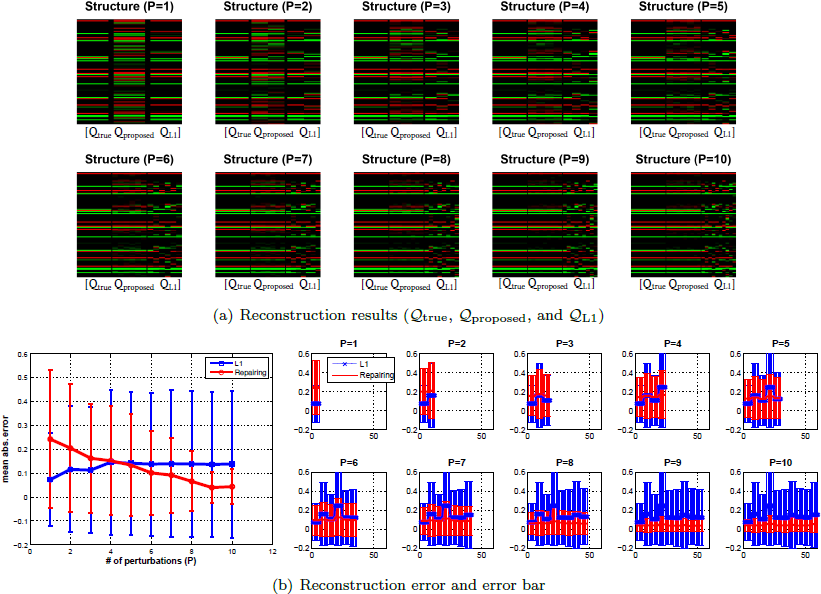
Reconstruction results and errors with respect to increasing the number of perturbation experiments *p*.

### Example 6

*(Convergence and Iteration) Since the algorithm iterates between the reconstruction and support estimation in Equation (11), we show the convergence results in Figure 8 where p* = 10. *Figure 8(left) shows the convergence of nuclear norm of* 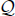 *with respect to the number of iterations; (right) shows convergence of reconstruction error and percentage of the number of recovered corruption region. At the first iteration, we only estimate a relatively small portion of corruption. However, as the number of iteration increases, we can refine the corrupted regions and identify a large amount of these regions.*

**Figure 8.**
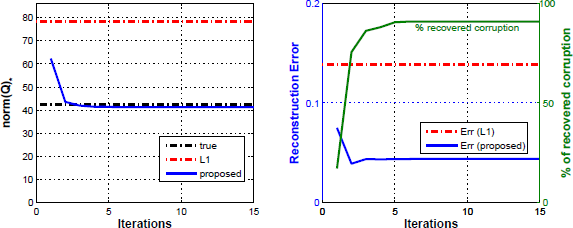
Example 6. Convergence with respect to increasing number of iterations *k*.

### Example 7

*(Nonlinear Dynamics: a large number of candidate basis functions) For a linear system, we choose the number of candidate basis functions (N* = *n) since we only consider simple linear relations. However, if we consider a nonlinear dynamical system, we increase the number of basis functions including simple linear term, monotonically increasing or decreasing Hill functions:*

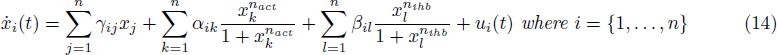

*where x*_*i*_ *denotes the concentration of the i-th species*, 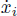 *is the change in concentration of the i-th species, the γ*_*ij*_ *denotes linear term, the α*_*ik*_, *β*_*il*_ *denote the maximum promoter/inhibitor strength, n*_*act*_ > 0 *represents positively cooperative binding (activation) and n*_*ihb*_ < 0 *represents negative cooperative binding (inhibition). First, we randomly generate GRN structure (i.e., γ*_*ij*_, *α*_*ik*_ and *β*_*il*_ *where we only consider diagonal term for γ*_*ij*_*, for example, protein decay rate) where n* = 10 *as shown in Figure 9(left) and simulate data with various perturbations (p* = 8 ≤ *n) as shown in Figure 9(right). Figure 10 presents both the reconstruction of the GRN and the estimated perturbation inputs with respect to increasing the number of sample time points. As the number of samples increases, the reconstructed GRN converges to the true GRN. However, the L*1 *reconstruction results only match the true graph structure partially since (unknown/unidentified) perturbation causes bias or uncertainties in the reconstruction result. Moreover, although we only allow 50% of the support information at the initial iteration, estimated inputs are consistent with true input in Figure 10(bottom).*

**Figure 9.**
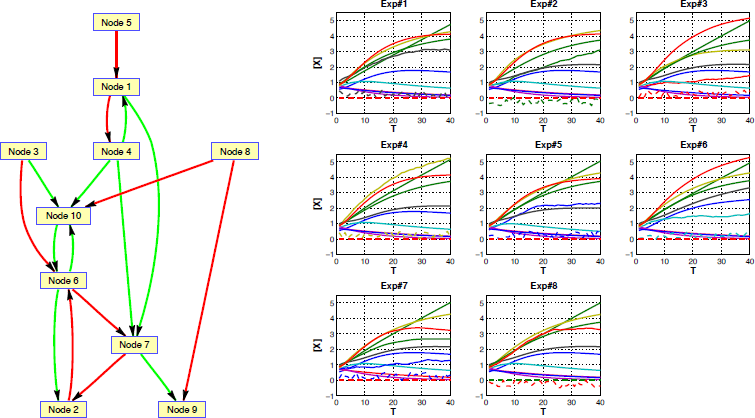
Example 7: Randomly generated GRNs (activation edges are shown in red, inhibition edges are shown in green and self-degradation loops (diagonal term of *γ*_*ij*_) are not presented here) and time series data set (solid line) with respect to various drug-induced perturbations (dotted line) where we use nonlinear ODEs and consider only one targeted drug for each experiment.

**Figure 10.**
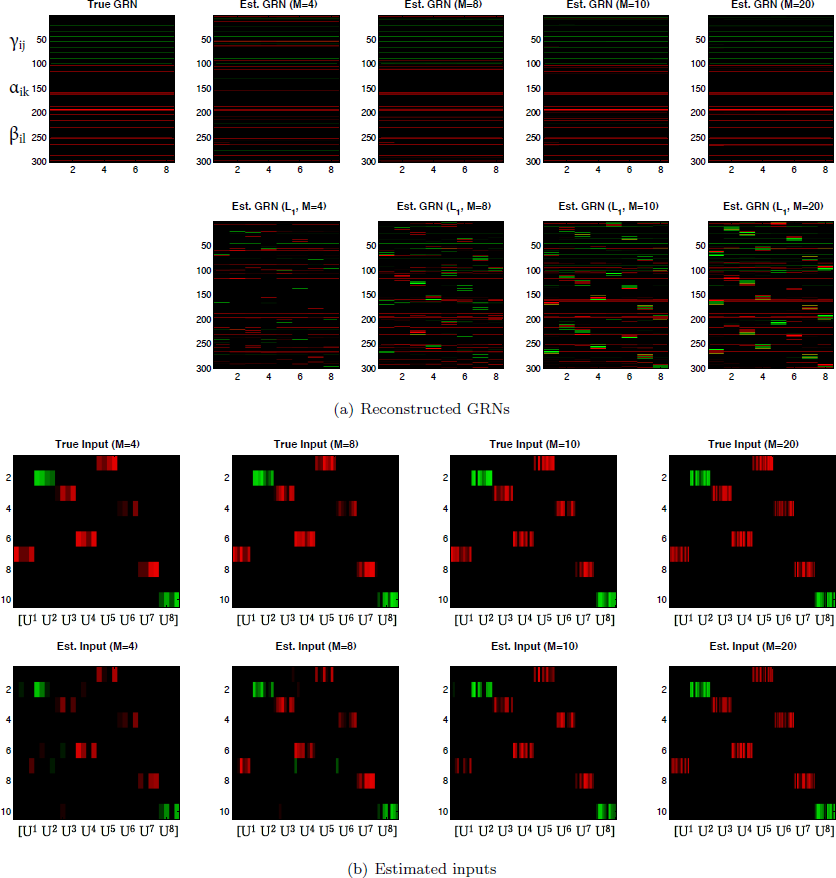
Example 7: Reconstructed GRNs and estimated input with respect to increasing the number of time steps (a) each row represents the corresponding *γ*_*ij*_ ∈ ℝ^100^, *α*_*ik*_ ∈ ℝ^100^, *β*_*il*_ ∈ ℝ^100^ and each column represents reconstructed GRN with respect to different experiments (*p* = 8) (b) each image represents 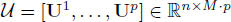 where **U**^*i*^ ∈ ℝ^*n*×*M*^. Here, we simply assume that each drug affects only the target protein (i.e., 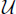 is partially block sparse; red color represents stimulation, green color represents inhibition and black color represents no perturbation in 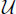) and add small sparse corruption.

## 6 Discussion

In this section, we demonstrate the practical relevance of the proposed method by applying it to the DREAM4 *in silica* Network Challenge dataset [28] [29] [30], a benchmark suite for performance evaluation of methods for gene network inference. Instead of random graph models, this dataset is generated by biologically plausible *in silica* networks, for example, by extracting sub-networks from transcriptional regulatory networks of *E. coli* and *S. cerevisiae* [28]. Also, time series datasets are generated from these networks using adequate dynamical models such as a detailed kinetic model of gene regulation. First, we show the result of GRN reconstruction based on the proposed method. Then, we interpret the implications of our result and use it to learn the relationship between the data and the identifiability of the proposed method.

### 6.1 Application of DREAM 4 *in silico* Network Challenge dataset

For the simplicity of analysis and explanation, we consider networks of size 10 and focus on their time series datasets with all perturbations. Each perturbation only affects about a third of all genes – but basal activation of these genes can be strongly increased or decreased. The genes that are directly targeted by the perturbation may then cause a change in the expression level of their downstream target genes leading to an indirect effect. As such, these experiments try to simulate physical or chemical perturbations applied to the cells, which would then cause some genes, via regulatory mechanisms, to have an increased or decreased basal activation.

The perturbations increase or decrease the basal activation of genes of the network simultaneously as shown in Figure 11(a). Each data set contains time courses showing how the GRN responds to a perturbation and how it relaxes upon removal of the perturbation. We consider 5 different time series and each time series has 21 time points (sampled every 50 steps). At *t* = 0, a perturbation is applied to the network, for example, a drug being added. The first half of the time series (until *t* = 500) shows the response of the network to the perturbation which is constantly applied from *t* = 0 to *t* = 500. At *t* = 500, the perturbation is removed and, thus, the second half of the time series (until *t* = 1000) shows how the gene expression levels go back from the perturbed to the unperturbed steady state. Since there are two different modes in time courses (i.e., with perturbation and without perturbation), in order to use all the time points, we should handle this perturbation condition properly. Otherwise, one model may not be able to fit both the first half and the second half of the time series.

**Figure 11.**
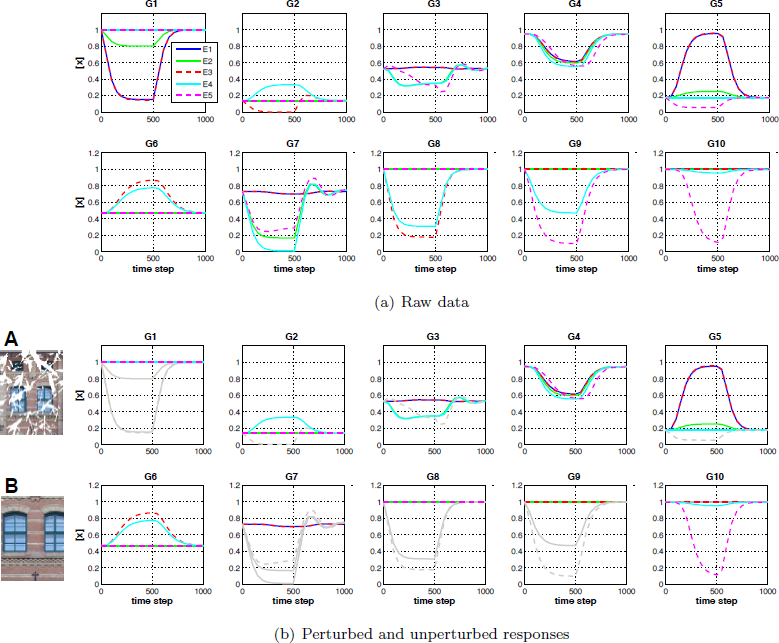
DREAM4 *in silico* dataset [28]: time series gene expressions are obtained by applying various perturbations to the original network in five different experiments. (a) raw data plots show dynamic responses of all genes across various perturbations (b) possible separation of the raw data based on the experiment design information shown in Table 1 where gray color denotes (possibly) corrupted responses by direct perturbations. One may consider the directly perturbed gene responses (gray color) as the corrupted image shown in **A** and the other responses (non-gray color) as the uncorrupted images shown in **B**.

**Table 1.**
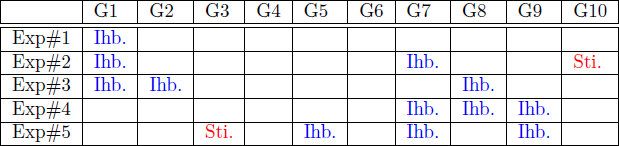
5 different experiment conditions where Ihb. represents inhibition and Sti. represents stimulation. This information can be used for initializing the support Ω_*i*_ in Equation (10) and separating perturbed and unperturbed responses shown in Figure 11(b).

|  | G1 | G2 | G3 | G4 | G5 | G6 | G7 | G8 | G9 | G10 |
| --- | --- | --- | --- | --- | --- | --- | --- | --- | --- | --- |
| Exp#1 | Ihb. |  |  |  |  |  |  |  |  |  |
| Exp#2 | Ihb. |  |  |  |  |  | Ihb. |  |  | Sti. |
| Exp#3 | Ihb. | Ihb. |  |  |  |  |  | Ihb. |  |  |
| Exp#4 |  |  |  |  |  |  | Ihb. | Ihb. | Ihb. |  |
| Exp#5 |  |  | Sti. |  | Ihb. |  | Ihb. |  | Ihb. |  |

Table 1 represents 5 different perturbation conditions, for example, for Exp#3 (the third row), Gene1 (G1), Gene2 (G2) and Gene8 (G8) are inhibited by drugs. Note that these are the known information, i.e., whether the response is corrupted by perturbation or not, but we do not know how much the perturbation affects the GRN response. Each treatment might also affect other genes, of which we have no *a priori* knowledge.

In order to infer the GRN structure using these time series gene expression data under various perturbations, we should identify how these perturbations affect a change in the expression level of the targeted genes. Otherwise, the inferred GRN can be biased or may only represent a partial structure of the whole GRN. To do this, we incorporate all data sets together and take advantage of the common structure of GRNs across the inferred GRNs. Since we only have partial information about the exact extent of the perturbations (or corruptions) as shown in Table 1, we should consider the (possibly) corrupted response as unmeasured and ignore it for reconstruction. By using the information in Table 1, we can initialize the support Ω_i_ in Equation (10) and further refine this support iteratively.

Figure 11(b) shows the initial separation of the (possibly) perturbed responses and the unperturbed responses based on the initial support. For example, from Table 1 and Figure 11(a), for Gene2 (G2), we know that we have to ignore Exp#3’s response which contains the (unknown) influence from perturbation (shown in gray color), and instead use the other experimental data (non-gray color) as shown in Figure 11(b). Similarly, for Gene3 (G3), we ignore Exp#5’s response but for Gene4 (G4), we can use all responses since there are no direct perturbations for G4. Intuitively, one may consider the directly perturbed gene responses (gray color) in Figure 11(b) as the analog of the corrupted parts of an image shown in Figure 11**A** and the other responses (non-gray color) as the analog of the uncorrupted parts of the image shown in Figure 11**B**.

Figure 12 shows the reconstruction result, the estimated corruption and the inferred GRN. In Figure 12(a)**A**, the first column represents the true GRN structure, where red represents activation and green represents inhibition edge. Since the proposed method uses both sparsity (which encourages sparsity of GRNs) and low-rank (which encourages commonalities across the inferred GRNs) of 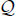, we can reconstruct the common GRN, most of which is consistent with the true GRN (the first column). Also, we can infer and estimate how much each perturbation affects the dynamics of the GRN, as depicted in Figure 12(a)**B**. Since we choose the set of possible candidate basis functions in Equation (6) and assume that the commonality is uniform across all genes and experiments, a small fraction of the reconstructed GRN is not consistent with the true GRN. We will further investigate and analyze this reconstruction result in the following section.

**Figure 12.**
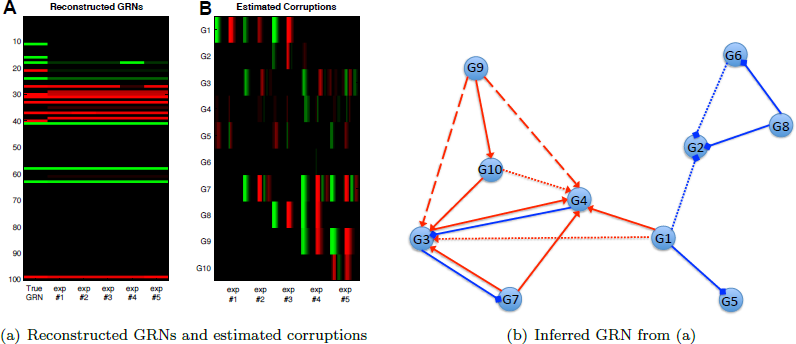
Reconstruction results: (a) reconstructed GRNs (**A**) where red color denotes the activation edge and green color denotes the inhibition edge and estimated corruption (**B**) where red color denotes positive values and green color denotes negative values. Note that the estimated corruptions represent temporal profiles which directly affect 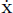, for example, when we perturb Gene1 (G1) in Exp#1, green color represents inhibition of G1 by drug perturbation; red color after green represents the effect of removing drug perturbation, (b) inferred GRN where solid lines denote the true positive (consistent with the true GRN), dotted lines denote the true negative (missing link) and dashed lines denote false positive (red: activation, blue: inhibition). Analyses and further details of these results are presented in Section 6.2.

### 6.2 Implications

In the previous section, we demonstrated the practical application to the DREAM4 data set. The proposed method is able to detect unknown influences caused by perturbations and then correctly repair the common graph structure across perturbed GRNs by isolating these effects in GRN inference.

In this section, we first analyze the reconstruction result together with the data set. Then, we discuss the implications of the proposed method through these analyses, discussing the relationship between the data and the identifiability of the network. We explain how one could optimize the experimental design to improve the identifiability of the network for the proposed method.

#### 6.2.1 Existence of *(dominant)* common dynamic responses

In order to estimate the effect of perturbations, the proposed method retrieves common dynamics of GRN subject to various perturbations. In other words, the proposed method uses low-rank of 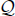 (or a combination of ‖·‖_1_ and total variation) to extract commonalities across different datasets. Thus, if these commonalities are not well exposed in the dataset, the method may fail to recover the corresponding components.

For instance, in Figure 12(b), the reconstruction result shows that we recover G1→G4 and G1⊣G5 but we fail to recover G1⊣G2 and G1→G3 (dotted line, missing link). In order to further investigate the reason for these results (true negative or missing link), we plot responses of G2, G3, G4 and G5 with respect to G1 in Figure 13. Again, the gray color denotes the response corrupted by perturbations. For example, in Figure 13**A**, G2 is directly perturbed in Exp#3 (gray color) and, thus, we ignore it for reconstruction.

**Figure 13.**
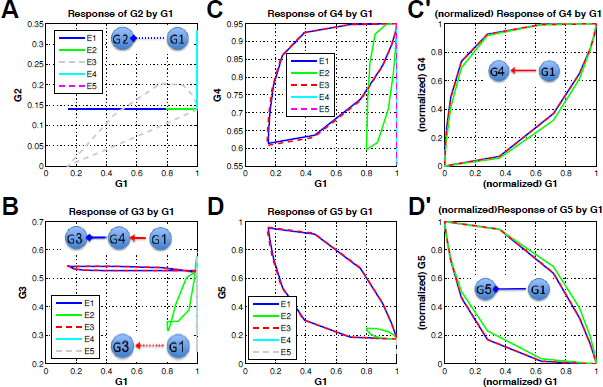
Responses of G2, G3, G4 and G5 with respect to G1 (influence from G1): each x-axis represents input response (i.e., G1) and each y-axis represents output responses (G2, G3, G4, and G5). (**A**) responses of G2 vs. G1 (**B**) responses of G3 vs. G1 (**C**) responses of G4 vs. G1 (**D**) responses of G5 vs. G1 (**C**′) (normalized) responses of G4 vs. G1 (**D**′) (normalized) responses of G5 vs. G1. Gray color represents the corrupted responses from perturbations which are ignored for reconstruction. For example, G2 (**A**) is directly perturbed in Exp#3, G3 (**B**) is directly perturbed in Exp#5 and G5 (**D**) is directly perturbed in Exp#5 as shown in Table 1.

- Figure 13**A**: Since the responses of G2 are not varying with respect to the responses of G1, there is no way to infer this connection (G1⊣G2) from this data set. Only the dataset for Exp#3 contains this inhibition response, however it also contains the unknown direct perturbation effect. Thus, the reconstruction result with this missing link is actually the best result for this dataset, as it avoids overfitting. Note that the response of G2 in Exp#4 (cyan color in Figure 13**A**) is not caused by G1 because G1 shows steady state response.
- Figure 13**B**: Given the data of Exp#1 and Exp#3, the response of G3 is more likely to be governed by (G1→G4⊣G3). Exp#2 is the only dataset that reflects the relationship (G1→G3). Thus, since the dynamic response corresponding to (G1→G3) is not a dominant common response for this dataset, we cannot reconstruct the corresponding GRN (G1→G3).
- Figure 13**C** and Figure 13**D**: Since all the responses show the consistency or the common dynamic responses, we can capture the true connections such as (G1→G4) and (G1⊣G5). Also, Figure 13**C** shows the effect of activation (positive correlation) and Figure 13**D** represents the effect of inhibition (negative correlation). For example, in Figure 13**C**, as G1 decreases, G4 decreases. On the other hand, in Figure 13**D**, as G1 decreases, G5 increases. In Figure 13**C**′ and Figure 13**D**′, we plot normalized responses to show the common dynamic responses clearly. Since dynamic features can be represented by possible candidate basis functions in Equation (6), the sparse low-rank representation can capture the commonality of the GRN structure.

This result implies that since the low-rank of 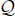 encourages commonalities across other’s GRNs, if there are no dominant common responses for a certain edge, it is challenging to infer the corresponding edge. In the context of image repairing, one can think that if a certain part of the sparse low-rank texture is not exposed well due to corruptions, we may not be able to repair such texture properly. For example, if the entire rows of the image in Figure 2(a) are corrupted, there is no way to correctly repair these rows.

Therefore, in order to reconstruct the GRN exactly, we have to design experiments with various perturbations that cause the underlying system to be perturbed and excited uniformly well. Or, we may have to consider different weighting factors across each gene for extracting the commonalities properly. For example, since we have more (uncorrupted) responses of G4 and G6, we can penalize the commonality more on G4 and G6. On the other hand, if there are only a few meaningful responses of a certain node, for example G3 in Figure 14**B**, we can reduce the weighting factor of commonality for that specific node.

Also, in practice, more information on drug perturbation (i.e., the GI-50 value) can help refine the effect of (unknown) perturbation. For example, in Figure 13**A**, if we know the effect of drug perturbation (i.e., independent dose-response data for drug), we can use the dataset for Exp#3 which contains the inhibition response (G1⊣G2) to infer the corresponding connection by isolating the effect of perturbation.

#### 6.2.2 Avoiding overfitting

As we discussed for Figure 13**A**, our method avoids overfitting and thus failed to infer the true inhibition (G1⊣G2) which could only be identified by corrupted data (gray color in Figure 13**A**). Similarly, in Figure 12(b), we reconstruct G8⊣G2 but fail to recover G6⊣G2. Figure 14**A** shows the response of G2 vs. G6. Since this response shows positive correlation and cannot match to inhibition (G6⊣G2), we fail to recover G6⊣G2. On the other hand, Figure 14**A**′ shows the response of G2 vs. G8 and matches to the effect of inhibition. Thus, we are able to capture this edge.

**Figure 14.**
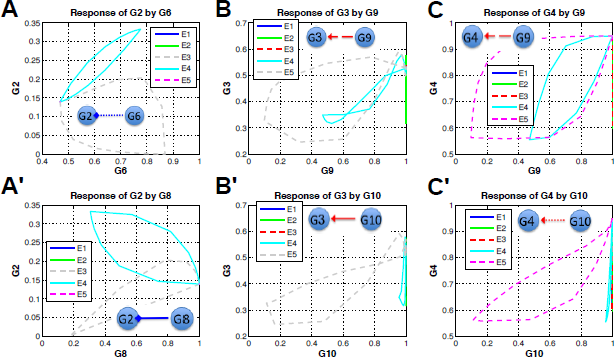
Input-output responses: each x-axis represents input response and each y-axis represents output response. (**A**/**A**′) responses of G2 with respect to G6 and G8 (**B**/**B**′) responses of G3 with respect to G9 and G10 (**C**/**C**′) responses of G4 with respect to G9 and G10. Gray color represents the responses corrupted by perturbations which are ignored for reconstruction.

Note that in Figure 14**A**′, the response of Exp#3 (gray color) shows both negative correlation (corresponding to true inhibition) and positive correlation (corresponding to direct perturbations). Since both G2 and G8 are inhibited by perturbations in Exp#3, if both perturbations’ effect are dominant, the response can show positive correlation. Thus, the responses driven by direct perturbations can distort the relationship between genes and lead to incorrect GRN inference. Therefore, if we cannot estimate and isolate the effect of these perturbations, we should ignore perturbed responses to avoid overfitting.

#### 6.2.3 Ambiguity

In Figure 12(b), the reconstruction results show false positive edges (dashed line, i.e., false discovery). Since the proposed method relies on the common dynamic features in the data sets to infer the GRN structure, if there are no dominant responses, it seems to be ambiguous and might present a challenge to infer the corresponding edge in the GRN structure. For example, consider influence at G3 and G4; since there is only one experiment which shows the dynamic response of G3 with respect to G9 in Figure 14**B** and G10 in Figure 14**B**′, it is hard to extract commonality. Thus, we identify a false positive link (G9→G3) (dashed line) in Figure 12(b). Similarly, since responses of G4 with respect to G9 show more common dynamic responses in Figure 14**C** and **D**, we infer a false positive link G9→G4 instead of G10→G4. Note that although the responses of G9 in Exp#4 and Exp#5 have similar scales in Figure 11(a) and G10 can only be affected by G9 (from the true GRN structure), the responses of G10 in Exp#4 and Exp#5 have quite different scales, which may cause a false positive inference.

#### 6.2.4 Validation

Since the proposed method can both extract common dynamic responses and estimate the effect of perturbations, we can also validate the reconstruction results by separating the dynamic responses into two sets; Figure 15 shows the separation of the dynamic responses into the common dynamic responses from GRN (**B**) and the aberration responses (**C**) where **A** represents the raw data, **B** represents dynamic responses from the inferred GRN and **C** represents the estimated corruptions. Note that the estimated corruptions shown in Figure 15**C** represents time course plots of Figure 12(a)**B** and these corruptions represent either large corruptions (i.e., the effect of drug perturbations) or small residuals between the raw data **A** and responses from sparse low-rank representation **B**. For instance, for G1 (the first column in Figure 15), since the responses are perturbed directly (Exp#1, Exp#2, Exp#3), the proposed method separates the response into the sparse error (corruption in **C**). However, for G4 (the fourth column in Figure 15), the sparse error represents small mismatch between the raw data **A** and the common responses from sparse low-rank representation **B**. Since we choose the set of possible candidate basis functions in Equation (6), there could be small mismatch between the raw data and the common responses from sparse low-rank representation.

**Figure 15.**
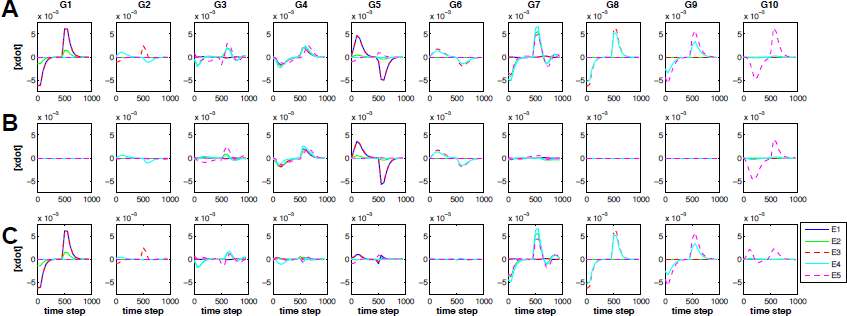
Separation of the dynamic responses: (**A**) the raw data, (**B**) the responses corresponding to the common GRN and (**C**) the estimated corruption including both the effects of perturbations and small mismatch between the raw data and the common responses from the inferred GRN.

Thus, we can validate the inferred GRNs with this information **C**, for example, the first column in Figure 12(b) (i.e., Exp#1) shows that large corruptions in G1 (drug perturbation) and small corruptions (residuals) in G4 and G5. Since we perturb G1 in Exp#1, we know that the large corruption in G1 represents drug effect and small corruption in G4 and G5 represents small mismatch between the raw data and the sparse low-rank representation (i.e., G1→G4 and G1⊣G5).

We now summarize the lessons learnt from the above analyses, where we applied our method to the DREAM data set. The proposed method

- can reconstruct the GRN by incorporating all data sets together.
- can incorporate from others’ inferred GRNs into the common GRN structure by using the low-rank property (*commonalities*).
- can infer and estimate the (*unknown*) drug effect, which can distort the relationship between genes and may lead to incorrect GRN inference in general, by separating the common dynamic response from the inferred GRN.
- can avoid overfitting but may fail to infer the true GRN when the dynamic responses corresponding to a certain edge do not show dominant common responses or they show ambiguities.

Since we develop the proposed method inspired by image repairing, these properties correspond to the underlying assumptions in image repairing, i.e., if there is no “Sparse Low-rank texture” representation in image, we cannot correctly repair the global structure of a corrupted texture. Therefore, in order to reconstruct the GRN exactly, all the genes in GRN should be perturbed uniformly well and any fractions of the responses should reflect the common responses enough for each gene.

## 7 Conclusion

In this paper, we show how to harness both sparse and low-rank structures for reconstructing GRNs in heterogeneous data sets based on various drug-induced perturbation experiments. Our method proposes a new convex formulation for GRN reconstruction and can automatically correctly repair the common graph structure of a partially perturbed GRN, even without precise information about the corrupting effects of drug-induced perturbations. Through synthetic experiment simulations and application of DREAM dataset, we show that our method can complete and repair GRN structure subjected to drug-induced perturbations. Also, through numerical comparisons, we demonstrate advantage over existing graph inference method dealing with different data sets and estimation of perturbation inputs. We are currently applying this method to large-scale datasets and using this tool for designing effective experiments in inferring the HER2+ breast cancer signaling pathway.

## Acknowledgments

This research was supported by the NIH NCI under the ICBP and PS-OC programs (5U54CA112970-08).

## Footnotes

1 Φ(*t*_*j*_) is known as the sensing matrix in compressive sensing [11]. Thus, for given sensing matrix Φ(*t*_*j*_) and measurement **y**(*t*_*j*_), we reconstruct s with penalizing sparsity (‖**s**‖_1_). In [11], we assume that **u**(*t*_*j*_) is known. Since we can simply subtract **u**(*t*_*j*_) from **y**(*t*_*j*_), we may reconstruct unbiased **s**. However, if **u**(*t*_*j*_) is not assumed to be known, this causes bias or uncertainties in reconstructing **s**.

